# Little evidence the standard genetic code is optimized for resource conservation

**DOI:** 10.1101/2021.02.04.429873

**Authors:** Hana Rozhoňová, Joshua L. Payne

## Abstract

Selection for resource conservation can shape the coding sequences of organisms living in nutrient-limited environments. Recently, it was proposed that selection for resource conservation, specifically for nitrogen and carbon content, has also shaped the structure of the standard genetic code, such that the missense mutations it allows tend to cause small increases in the number of nitrogen and carbon atoms in amino acids. Moreover, it was proposed that this optimization is not confounded by known optimizations of the standard genetic code, such as for polar requirement or hydropathy. We challenge these claims. We show the proposed optimization for nitrogen conservation is highly sensitive to choice of null model and the proposed optimization for carbon conservation is confounded by the known conservative nature of the standard genetic code with respect to the molecular volume of amino acids. There is therefore little evidence the standard genetic code is optimized for resource conservation.

---

The standard genetic code (SGC) exhibits numerous optimizations (Freeland et al. 2003). For example, the missense mutations it allows tend to preserve key physicochemical properties of amino acids, such as polar requirement, hydropathy (Haig and Hurst 1991), and to a lesser extent, molecular volume (Haig and Hurst 1999). Recently, an additional optimization was proposed, namely for resource conservation (Shenhav and Zeevi 2020). Motivated by the observation that selection for resource conservation can shape the coding sequences of organisms living in nutrient-limited environments (Mazel and Marlière 1989; Elser et al. 2006; Bragg and Wagner 2007; Lv et al. 2008; Li et al. 2009; Grzymski and Dussaq 2012; Mende et al. 2017; Hellweger et al. 2018), it was hypothesized that selection for resource conservation has also shaped the structure of the SGC, such that the missense mutations it allows tend to cause small increases in the number of nitrogen and carbon atoms in amino acids. Moreover, it was hypothesized that this optimization is not confounded by known optimizations of the SGC, such as for polar requirement or hydropathy. These hypotheses were tested by quantifying the expected random mutation cost (ERMC) of missense mutations allowed by the SGC, measured in units such as the number of nitrogen or carbon atoms or the absolute change in polar requirement or hydropathy of amino acids, and comparing this cost to those incurred by a large number of alternative codes (Shenhav and Zeevi 2020).

Drawing comparisons with alternative codes has a long history in the study of optimizations in the SGC (Alff-Steinberger 1969; Haig and Hurst 1991). Because the space of alternative codes is so large (≈ 1.5 · 10^84^) (Caporaso et al. 2005), it is necessary to draw comparisons with a sample from this space, which can then be used as a null model (Freeland et al. 2003). There are many methods for generating alternative codes, and different methods can generate codes with different properties (Wichmann and Ardern 2019). For example, a method known as quartet shuffling generates alternative codes by randomly shuffling quartet blocks — the blocks of four codons that share the first two bases (e.g., AAA, AAC, AAG, AAU) (Alff-Steinberger 1969; Caporaso et al. 2005). This method, which was used to test the hypothesis that the SGC is optimized for resource conservation (Shenhav and Zeevi 2020), generates alternative codes that preserve two key properties of the SGC, namely the number of codons per amino acid and the degeneracy of the third base (e.g., the three codons for isoleucine always have the same first and second base as the codon for methionine). In contrast, a method known as amino acid permutation generates alternative codes by randomly permuting the twenty standard amino acids amongst the synonymous codon blocks (Haig and Hurst 1991). This method, which is most commonly used in the field (Haig and Hurst 1991; Ardell 1998; Freeland and Hurst 1998; Freeland et al. 2000; Gilis et al. 2001; Archetti 2004; Caporaso et al. 2005; Goodarzi et al. 2005a; Goodarzi et al. 2005b; Novozhilov et al. 2007; Butler et al. 2009; Tripathi and Deem 2018), generates alternative codes that preserve a different key property of the SGC, namely the structure of the synonymous codons blocks. Importantly, in these alternative codes, the number of codons per amino acid can change drastically relative to the SGC, because the permutation of amino acids amongst the synonymous blocks is random. These two methods therefore generate alternative codes with substantially different structural properties (Fig. S1).

Here, we test the hypothesis that the SGC is optimized for resource conservation with respect to nitrogen and carbon content by drawing comparisons with alternative codes generated using ten different methods, including quartet shuffling and amino acid permutation (Table 1; Supplementary Methods). With respect to nitrogen, we find consistent statistical support for resource conservation using only one of the ten methods. With respect to carbon, we find consistent statistical support for resource conservation across the ten methods, but show that this optimization is confounded by the known conservative nature of the SGC with respect to the molecular volume of amino acids (Haig and Hurst 1999).

**Table 1:** Two key properties of the alternative genetic codes generated with the ten different methods used in this study (Supplementary Methods).

| Method | Preserves the number of codons per amino acid | Preserves the exact block structure of the SGC |
| --- | --- | --- |
| *Quartet shuffling* | yes | no<sup>a</sup> |
| *Amino acid permutation* | no | yes |
| *Restricted amino acid permutation* | no<sup>b</sup> | yes |
| *N-Block Shuffler* | yes | no<sup>a</sup> |
| *Codon Shuffler* | yes | no |
| *AAAGALOC Shuffler* | no | no |
| *Random expansion* | no | yes |
| *Ambiguity reduction 1* | no | yes |
| *Ambiguity reduction 2* | no | yes |
| *2-1-3 model* | no | yes |

<sup>a</sup> The alternative codes have a block structure, but it is different from that of the SGC.
<sup>b</sup> The number of codons per amino acid is allowed to change by at most two, relative to the SGC.

## Results

### Computing the expected random mutation cost

We compute the expected random mutation cost (ERMC) as

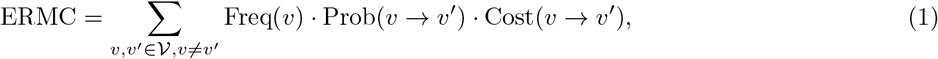

where *ν* is the set of all 64 codons, Freq(*υ*) is the frequency of codon *υ* (Supplementary Note 1), Prob(*υ* → *υ*′) is the probability of mutation from codon *υ* to *υ*′ given a genetic code (standard or otherwise) and mutation rates (e.g., based on a transition:transversion ratio), and Cost(*υ* → *υ*′) is the cost of mutating codon *υ* to *υ′* (Shenhav and Zeevi 2020). For resource conservation, the cost is defined as the increase in the number of nitrogen or carbon atoms in the amino acid encoded by codon *υ*′ relative to the amino acid encoded by *υ*, whereas for amino acid properties such as polar requirement, hydropathy, and molecular volume, it is defined as the absolute difference in the respective property of the amino acid encoded by codon *υ*′ and the amino acid encoded by *υ* (Shenhav and Zeevi 2020).

We use three sets of codon frequencies and mutation rates to compute the ERMC of a genetic code (Shenhav and Zeevi 2020) (Supplementary Methods), namely

1. *Baseline parameters*: All codon frequencies are equal and mutation rates are based on a transition:transversion ratio of 1:2.
2. *Ocean parameters*: Codon frequencies and mutation rates are derived from marine metagenomics samples (Shenhav and Zeevi 2020).
3. *Diverse species parameters* : Codon frequencies are derived from 39 species (Athey et al. 2017) and mutation rates are based on 11 transition:transversion ratios ranging from 1:5 to 5:1. In total, this set includes 429 combinations of codon frequencies and mutation rates (Shenhav and Zeevi 2020).

For each set, we determine the statistical significance of the ERMC of the SGC by computing an empirical *p*-value, which is the fraction of 1 million alternative genetic codes that have an ERMC that is less than or equal to that of the SGC. We compute this empirical *p*-value separately for each of the ten methods for generating alternative codes. For the *diverse species parameters*, we correct the *p*-values for testing multiple hypotheses (Benjamini and Hochberg 1995). We report the raw and corrected *p*-values for all tests in Supplementary Data S1-S9.

### Nitrogen conservation is highly sensitive to choice of null model

We find consistent statistical support for nitrogen conservation in the SGC using only one of the ten methods for generating alternative codes (Supplementary Data S1), namely the codon shuffler (Caporaso et al. 2005) (*Baseline parameters*: *p* = 1.00·10^−6^; *Ocean parameters*: *p* = 3.00 · 10^−6^; *Diverse species parameters*: *p* ≤ 0.016). For the remaining nine methods, nitrogen conservation is never consistently statistically significant across all tested parameters, as illustrated for amino acid permutation in Fig. 1 (*Baseline parameters*: *p* = 0.485; *Ocean parameters*: *p* = 0.115; *Diverse species parameters* : *p* ≥ 0.573). Surprisingly, even for alternative codes generated by quartet shuffling, nitrogen conservation is not statistically significant for the *diverse species parameters* (*p* ≥ 0.316), although it is for the *baseline parameters* (*p* = 0.023) and *ocean parameters* (*p* = 0.034).

**Figure 1:**
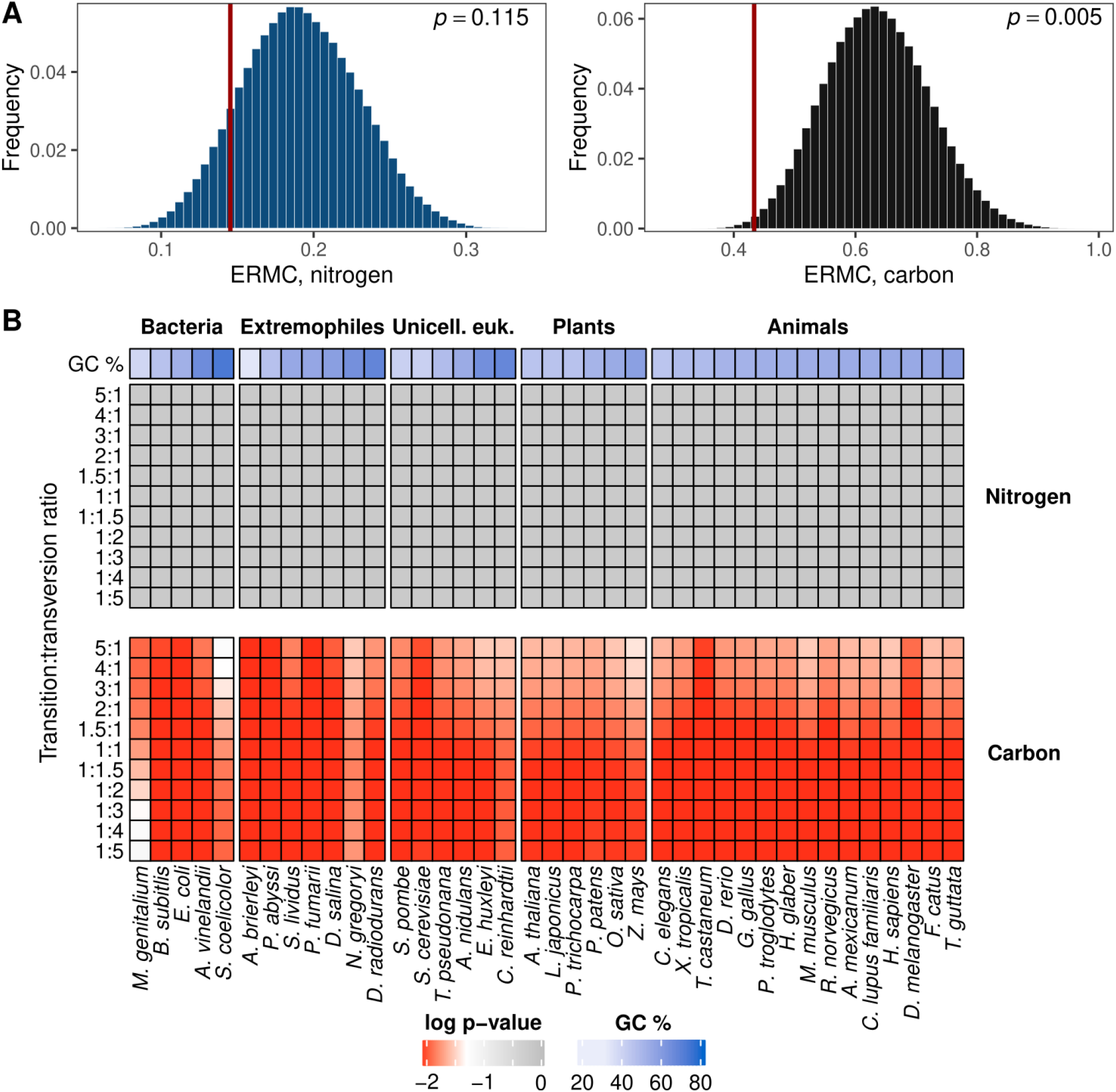
Nitrogen conservation is highly sensitive to choice of null model.

A – Histograms of the ERMC for nitrogen (blue) and carbon (black) in 1 million alternative codes generated by amino acid permutation. The vertical red line corresponds to the SGC. Codon frequencies and mutation rates are from the *ocean parameters*.
B – P-values of the ERMC for nitrogen (top) and carbon (bottom) of the SGC, relative to 1 million alternative codes generated by amino acid permutation, using the *diverse species parameters*. Shades of gray correspond to statistically insignificant *p*-values (*p* > 0.05; darker = less significant) and shades of red to statistically significant *p*-values (*p ≤* 0.05; darker = more significant). The *p*-values were adjusted using Benjamini-Hochberg correction for multiple testing. Organisms in each group are ordered based on the GC content of their coding sequences. Unicell. euk. = unicellular eukaryotes.

What explains the qualitative difference between the results obtained using these different methods? The key is that the alternative codes generated by both quartet shuffling and the codon shuffler maintain the number of codons per amino acid. In the SGC, the codons of the six nitrogen-rich amino acids (i.e., those with at least one nitrogen atom in their side chain: histidine, lysine, asparagine, glutamine, arginine, and tryptophan) are clustered in the codon table (Fig. S1A; Supplementary Note 2), such that a point mutation to a codon of a nitrogen-rich amino acid leads with probability 0.489 to a codon of the same or different nitrogen-rich amino acid. Because such clustering is highly unlikely in alternative codes that maintain the number of codons per amino acid, these almost always have a higher ERMC for nitrogen than the SGC, thus rendering nitrogen conservation statistically significant. In contrast, if the number of codons per amino acid is allowed to change, many alternative codes have a lower ERMC for nitrogen than the SGC. The reason is that the ERMC for nitrogen is strongly correlated with the number of codons for nitrogen-rich amino acids in these alternative genetic codes (Pearson’s correlation 0.567, *p* < 2.2 · 10^−16^ for codes generated by amino acid permutation; Figure 2) and this number is often smaller than in the SGC, thus rendering nitrogen conservation statistically insignificant.

**Figure 2:**
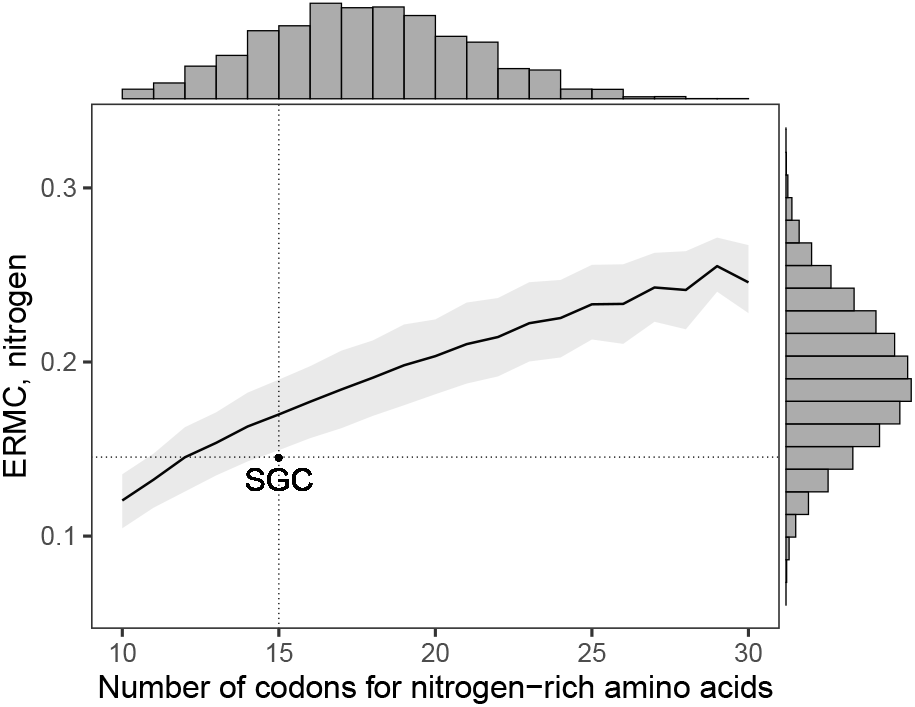
The ERMC for nitrogen is correlated with the number of codons for nitrogen-rich amino acids. The black line shows the mean, and the shaded area shows the 25th to the 75th quantile, of the ERMC for nitrogen in relation to the number of codons for nitrogen-rich amino acids in 1 million alternative codes generated by amino acid permutation. The point and dotted lines correspond to the SGC. Histograms of the number of codons for nitrogen-rich amino acids and the ERMC for nitrogen are shown on the top and on the right of the main panel, respectively. The ERMC for nitrogen was computed using the *ocean parameters*.

### Carbon conservation is confounded by the molecular volume of amino acids

We find consistent statistical support for carbon conservation in the SGC across the ten methods for generating alternative codes (*Baseline parameters*: *p* < 0.05 for 10 of 10 methods; *Ocean parameters*: *p* < 0.05 for 9 of 10 methods; *Diverse species parameters*: median *p* < 0.05 for 9 of 10 methods; Fig. 1; Supplementary Data S2). However, we hypothesize that carbon conservation is confounded by molecular volume (Grantham 1974), because the molecular volume of an amino acid is strongly correlated with its number of carbon atoms (Pearson’s correlation 0.906, *p* = 3.97 · 10^−8^; Fig. 3A) and the changes caused by missense mutations to amino acids’ molecular volume and number of carbon atoms are therefore strongly correlated (Pearson’s correlation 0.813, *p* < 2.2 · 10^−16^; Fig 3B). We test this hypothesis using a hierarchical model (Shenhav and Zeevi 2020). Specifically, for each of the ten methods for generating alternative codes, we examine the subset of alternative codes that have an ERMC that is less than or equal to that of the SGC for molecular volume and test whether the SGC is also optimized for carbon conservation relative to this subset. It is not (*Baseline parameters*: *p* > 0.05 for 10 of 10 methods; *Ocean parameters*: *p* > 0.05 for 10 of 10 methods; *Diverse species parameters*: minimum *p* > 0.05 for 9 of 10 methods; Supplementary Data S3), as illustrated in Fig. 3C for alternative codes generated by amino acid permutation (*Baseline parameters*: *p* = 0.139; *Ocean parameters*: *p* = 0.125; *Diverse species parameters*: *p* > 0.190). Thus, carbon conservation is confounded by the known conservative nature of the SGC with respect to the molecular volume of amino acids (Haig and Hurst 1999).

**Figure 3:**
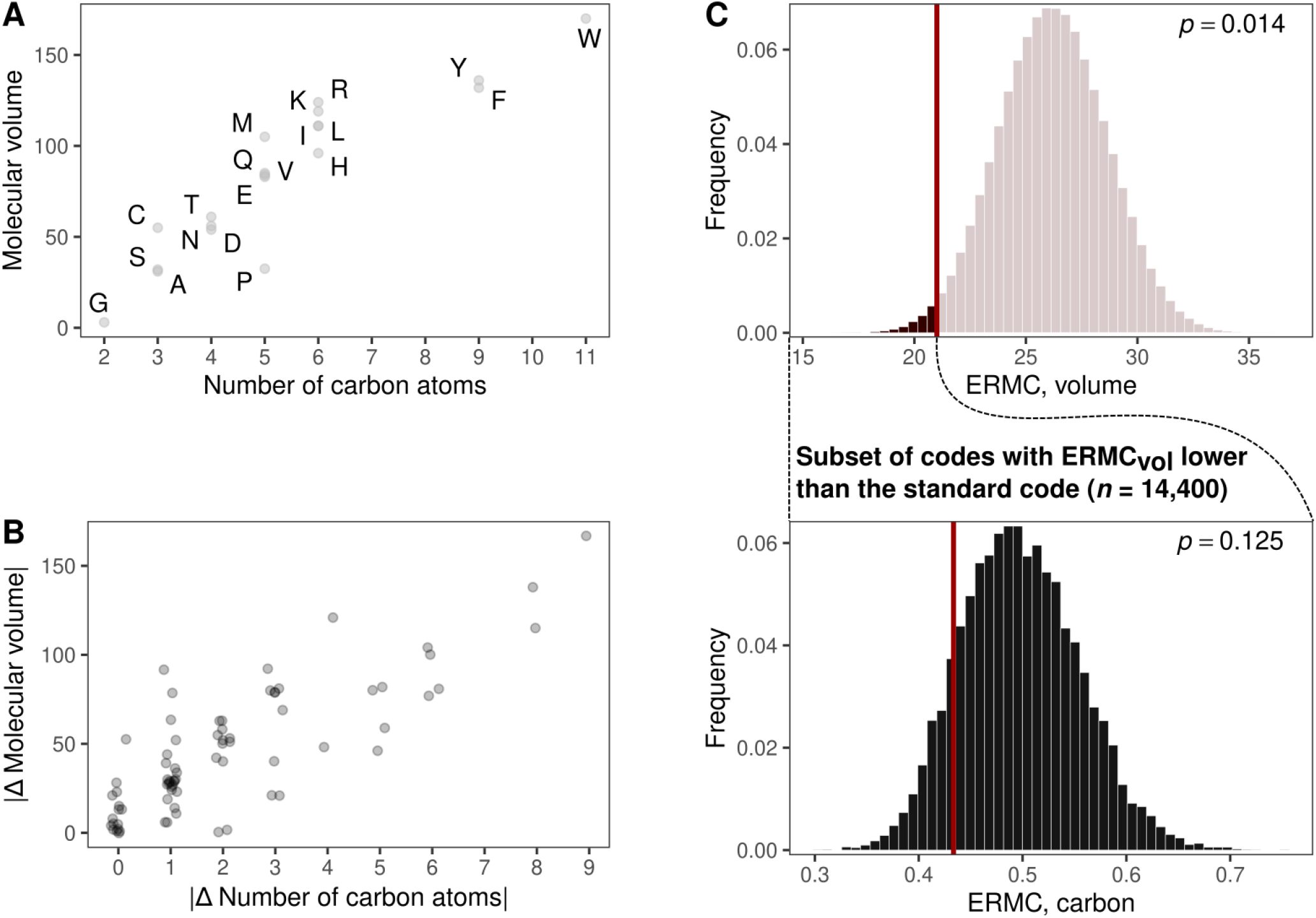
Carbon conservation is confounded by the molecular volume of amino acids.

A – Scatter plot of the number of carbon atoms and the molecular volume of the twenty proteinogenic amino acids.
B – Scatter plot of the absolute change in the number of carbon atoms and the absolute change in molecular volume for the 75 amino acid pairs that are connected by a missense mutation in the SGC. Jitter applied in the x-axis for visualization.
C – Histograms of the ERMC for (top) the molecular volume of amino acids in 1 million alternative codes generated by amino acid permutation and (bottom) for carbon in the subset of 14,400 alternative codes that have an ERMC for molecular volume that is less than or equal to that of the SGC. The ERMC was computed using the *ocean parameters*.

## Discussion

We found that the proposed optimization of the SGC for nitrogen conservation (Shenhav and Zeevi 2020) is highly sensitive to choice of null model. Specifically, we only found statistical support for nitrogen conservation when using null models that preserve the number of codons per amino acid from the SGC. Choosing an appropriate null model to test for optimizations in the SGC is challenging, because different null models preserve different key properties of the SGC, while perturbing others. Which key properties should be preserved and which should be perturbed? This is a difficult question, but our knowledge of extant non-standard genetic codes, which are derived from the SGC (Knight et al. 2001), clearly indicate that the number of codons per amino acid is not one of these key properties. The same can be said for the structure of the synonymous codon blocks. Given this challenge, a sensible way forward is to use a diversity of null models when testing for optimizations in the SGC (Wichmann and Ardern 2019) and to refrain from reporting optimizations that only find statistical support from a small number of these null models.

Indeed, we found such broad statistical support across a diversity of null models for the proposed optimization for carbon conservation (Shenhav and Zeevi 2020), but we also found that this optimization is confounded by the known conservative nature of the SGC with respect to molecular volume (Haig and Hurst 1999). This highlights another challenge in choosing an appropriate null model to test for optimizations in the SGC: Most null models are agnostic to the evolutionary history of the SGC, which can give the false impression that optimizations are the product of selection rather than a byproduct of the physical processes of gene duplication and mutation (Stoltzfus and Yampolsky 2007; Massey 2008). Carbon conservation is a case in point. A prominent model of the evolutionary history of the SGC suggests that in the early stages of code evolution, amino acids were recognized by pockets in the tertiary structure of proto-tRNAs and that the expansion of the code proceeded via duplication and mutation of these proto-tRNAs (Wolf and Koonin 2007). Because a recently duplicated proto-tRNA would likely recognize an amino acid with similar molecular volume to that recognized by its parent proto-tRNA (Massey 2008), gene duplication and mutation would naturally result in a clustering of codons for amino acids with similar molecular volumes in the codon table, as present in the SGC. As carbon is the main building block of all proteinogenic amino acids, the proposed optimization for carbon conservation follows naturally, without needing to evoke selection for resource conservation.

Finally, we note that if in nutrient-limited environments it is costly for missense mutations to increase the number of nitrogen or carbon atoms in amino acids, then it should be beneficial for missense mutations to decrease the number of nitrogen or carbon atoms in amino acids. This simple fact makes it difficult to justify the ERMC as a measure of the cost of missense mutations, because it only accounts for increases. Indeed, contemporaneous work to ours shows that the SGC is not optimized for resource conservation when the ERMC is modified to also account for decreases in the number of nitrogen or carbon atoms (Xu and Zhang 2021). Taken together, our analyses strongly suggest that the SGC is not optimized for resource conservation.

## Acknowledgements

We thank Sinisa Bratulic and David M. McCandlish for discussions and feedback on this manuscript. This work was supported by the Swiss National Science Foundation (grant PP00P3 170604).

## Data availability

Code used in this study is freely available at https://github.com/parizkh/resource-conservation-in-genetic-code.

## Supplementary methods

### Generating alternative codes

We use ten diverse methods for generating alternative genetic codes:

- *Quartet shuffling* (Alff-Steinberger 1969; Caporaso et al. 2005; Shenhav and Zeevi 2020): This method randomly permutes the so-called quartet blocks — quartets of codons that share the first and second nucleotide (e.g., AAA, AAC, AAG, and AAU). Moreover, it requires that the two sets of stop codons are separated by a single transition mutation (Shenhav and Zeevi 2020).
- *Amino acid permutation* (Haig and Hurst 1991): This method randomly permutes the twenty amino acids amongst the synonymous codon blocks.
- *Restricted amino acid permutation*: This method is the same as amino acid permutation, except the set of permutations is restricted to those in which the number of codons per amino acid changes by at most two, relative to the SGC.
- *N-Block Shuffler* (Caporaso et al. 2005): This method randomly permutes amino acids amongst synonymous codon blocks of the same size.
- *Codon Shuffler* (Caporaso et al. 2005): This method randomly assigns codons to amino acids, ensuring that the number of codons per amino acid is the same as in the SGC.
- *AAAGALOC Shuffler* (Caporaso et al. 2005): This method generates alternative codes at random, only requiring that each of the twenty amino acids has at least one codon.
- *Random expansion* (Massey 2008): This method generates alternative codes by the sequential assignment of amino acids to codon blocks. The first amino acid is chosen at random and is assigned to a randomly chosen codon block. Subsequent amino acids are also chosen at random, but with probability proportional to the inverse of their Grantham distance (Grantham 1974) to the last amino acid added to the code^1^. This amino acid is assigned at random to one of the codon blocks that have at least one codon within one mutation from a codon for the last amino acid added to the code. This method is called ‘Model 1’ in Massey (2008).
- *Ambiguity reduction 1* (Massey 2008): Similarly to *Random expansion*, this method generates alternative codes by sequential assignment of amino acids to codon blocks. Initially, there are only two large codon blocks, each consisting of 32 codons. The amino acids encoded by these two codon blocks are chosen at random. These large codon blocks are then subsequently split into smaller blocks, in a specified order. In each step, a codon block is split into two smaller blocks; one of them encodes the amino acid originally encoded by the block before splitting, while the second one is assigned a randomly chosen amino acid, chosen with probability proportional to the inverse of its Grantham distance to the original amino acid. This method is called ‘Model 2a’ in Massey (2008).
- *Ambiguity reduction 2* (Massey 2008): This method works the same as *Ambiguity reduction 1*, only the order in which the codon blocks are split is different. It is called ‘Model 2b’ in Massey (2008).
- *2-1-3 model* (Massey 2008): Similar to *Ambiguity reduction 1* and *Ambiguity reduction 2*, this method generates alternative codes by splitting the codon blocks according to the 2-1-3 model, i.e., assuming that the codon positions acquired coding potential in the order of the second, first, and then third position (Massey 2006). The first four amino acids are fixed (V, A, D, G, encoded by codons with U, C, A, and G, respectively, in the second codon position). This method is called ‘Model 3’ in Massey (2008).

For all methods except quartet shuffling, the positions of the stop codons were fixed as in the SGC.

### Codon frequencies and mutation rates

For the *ocean parameters*, we obtained the codon frequencies and mutation rates directly from David Zeevi (Shenhav and Zeevi 2020). For the *diverse species parameters*, we obtained the codon frequencies from the source code of Shenhav and Zeevi (2020).

### Correcting for multiple testing

For the *diverse species parameters*, the *p*-values were corrected using the `p.adjust` function in R, with the method parameter equal to `“BH”` (Benjamini-Hochberg correction for multiple testing (Benjamini and Hochberg 1995)).

## Supplementary Note 1

The ERMC accounts for codon frequencies (Eq. 1), which are known for many species (Athey et al. 2017). As originally formulated, the ERMC does this accounting by setting Freq(*υ*) to the median frequency of all codons for amino acid *a*, rather than using the actual frequency of codon *υ* from the species of interest (Shenhav and Zeevi 2020). That is, it assumes that all codons for the same amino acid have the same frequency. What is more, this median frequency is always calculated using the SGC, even when measuring the ERMC of alternative codes. That is, the frequency of codons encoding amino acid *a* is always the same as in the SGC, no matter which codons map to amino acid *a* in an alternative code. While an argument could be made for assuming a fixed abundance of amino acids in the proteome, we do not see the biological relevance of extending this argument to assuming fixed codon frequencies. In making it, valuable information on codon frequencies is lost, which is significant because codon usage patterns differ widely among species (Plotkin and Kudla 2011), also due to selection for resource conservation (Shenhav and Zeevi 2020). We therefore do not follow this treatment of codon frequencies in our main analyses, and instead use codon frequencies directly as they were reported for each species of interest.

Nonetheless, to facilitate direct comparison to previous work (Shenhav and Zeevi 2020), we repeat all of our analyses using this treatment of codon frequencies. Defining the abundance of amino acid *a* as the sum of the frequencies of all codons encoding *a* in the SGC, we consider two alternative definitions of the frequency Freq(*υ*) of codon *υ* in an alternative code:

1. *Median frequencies*: Freq(*υ*) is defined as the median frequency of all codons encoding the same amino acid as *υ* in the SGC. This treatment is identical to how the codon frequencies were defined by Shenhav and Zeevi (2020). However, if the number of codons per amino acid is allowed to change in the alternative codes, this method does not preserve the abundance of each amino acid from the SGC in the alternative codes.
2. *Mean frequencies*: Freq(*υ*) is defined as the abundance of the amino acid encoded by *υ* divided by the number of codons encoding the amino acid in the alternative code. This treatment is different from how the codon frequencies were defined by Shenhav and Zeevi (2020), because it takes the mean rather than the median. However, if the number of codons per amino acid is allowed to change in the alternative codes, this method preserves the abundance of each amino acid from the SGC in the alternative codes.

Our results are qualitatively unchanged by these modified codon frequencies, with two exceptions: (1) Nitrogen conservation becomes statistically significant for one additional method for generating alternative codes, namely the N-Block shuffler, for both alternative treatments of codon frequencies (*Median frequencies*: *Baseline parameters*: *p* = 0.015; *Ocean parameters*: *p* = 3.88 · 10^−3^; *Diverse species parameters*: median *p* = 0.032; *Mean frequencies*: *Baseline parameters*: *p* = 0.015; *Ocean parameters*: *p* = 3.32 · 10^−3^; *Diverse species parameters*: *p* ≤ 0.047). (2) Carbon conservation becomes statistically significant after controlling for the confounding factor of molecular volume for 4 of 10 methods for generating alternative codes, but only when using the *ocean parameters* and *mean frequencies* (Supplementary Data S4-S9; *p*-values computed using a set of 100,000 alternative codes).

## Supplementary Note 2

It was proposed that nitrogen conservation is driven by structural principles of the SGC, namely by the arrangement of synonymous codon blocks for nitrogen-rich amino acids in the codon table (Shenhav and Zeevi 2020). The SGC exhibits a ‘square arrangement’, in which codons of five of the six nitrogen-rich amino acids (histidine, lysine, asparagine, glutamine, and arginine, but not tryptophan) span only two nucleotides in their first and second positions (codons CAN, CGN, AAN and AGR, with N denoting any nucleotide and R denoting A or G). Using quartet shuffling to generate alternative codes, it was argued that this ‘square arrangement’ (Figure S2A, middle) amplifies nitrogen conservation relative to other arrangements, such as a ‘diagonal arrangement’, in which the four codon blocks span all four nucleotides in both their first and second positions (Figure S2A, right) (Shenhav and Zeevi 2020). We uncover an additional arrangement that is even more conducive to nitrogen conservation, which we call a ‘line arrangement’ (Figure S2A, left). In it, all codons for nitrogen-rich amino acids share the same nucleotide in the first or the second position (i.e., these codons occupy one row or one column of the codon table). When generating alternative codes with quartet shuffling, those that exhibit a ‘line arrangement’ tend to have a significantly lower ERMC for nitrogen than alternative codes with other arrangements (Fig. S2B; e.g., *p* < 2.2 · 10^−16^, ‘line arrangement’ vs. ‘square arrangement,’ Welch two sample t-test). The reason is the ‘line arrangement’ maximizes the number of missense mutations between codons of nitrogen-rich amino acids, and hence minimizes the number of missense mutations from codons for non-nitrogen-rich amino acids to codons for nitrogen-rich amino acids.

---

**Figure S1:**
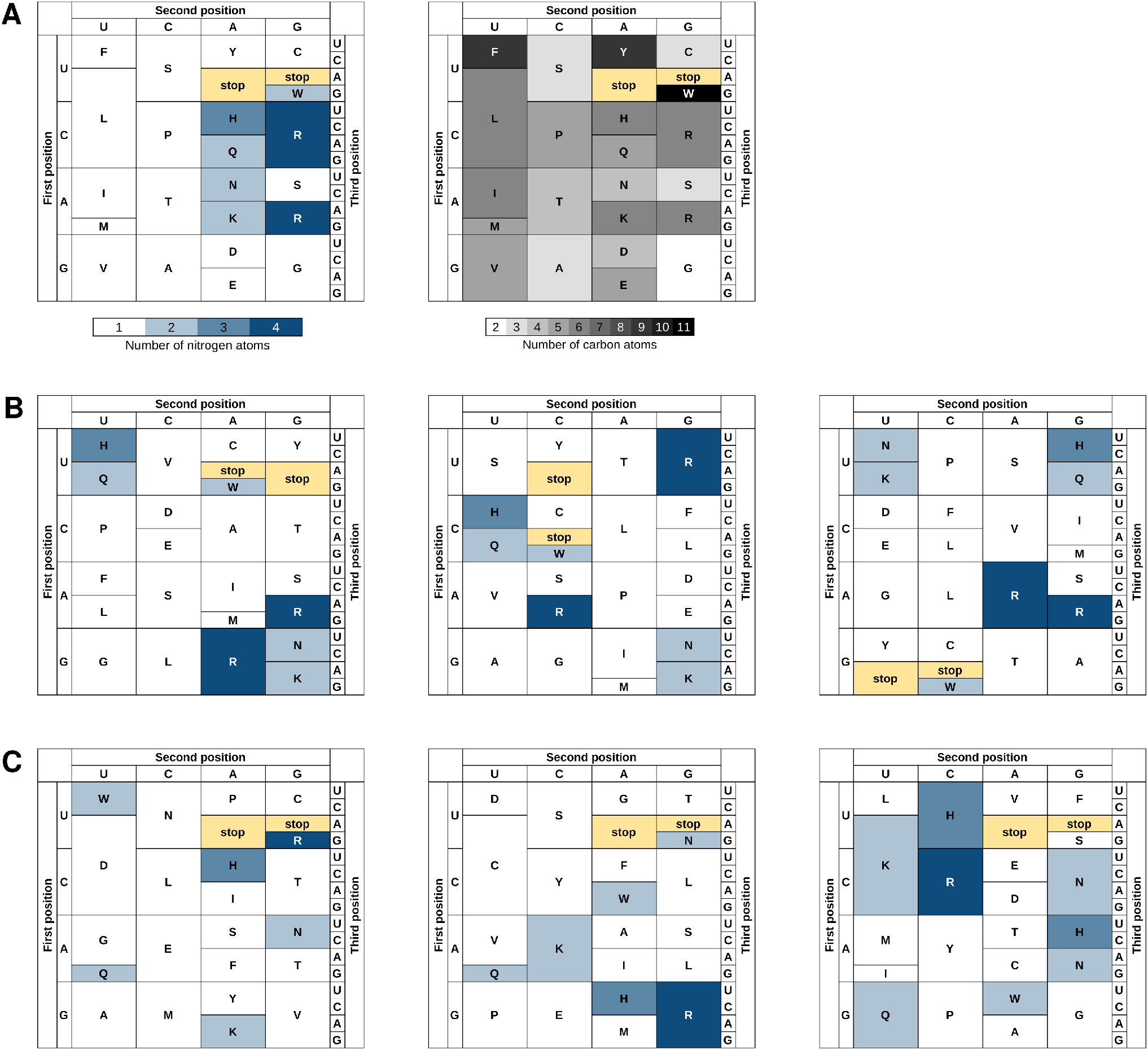
The standard genetic code and examples of alternative codes.

A – The standard genetic code. The codons are colored based on the (left) nitrogen and (right) carbon content of the corresponding amino acid.
B – Three examples of codes generated by quartet shuffling (Alff-Steinberger 1969; Caporaso et al. 2005; Shenhav and Zeevi 2020).
C – Three examples of codes generated by amino acid permutation (Haig and Hurst 1991). In B and C, the codons are colored based on the nitrogen content of the corresponding amino acid.

**Figure S2:**
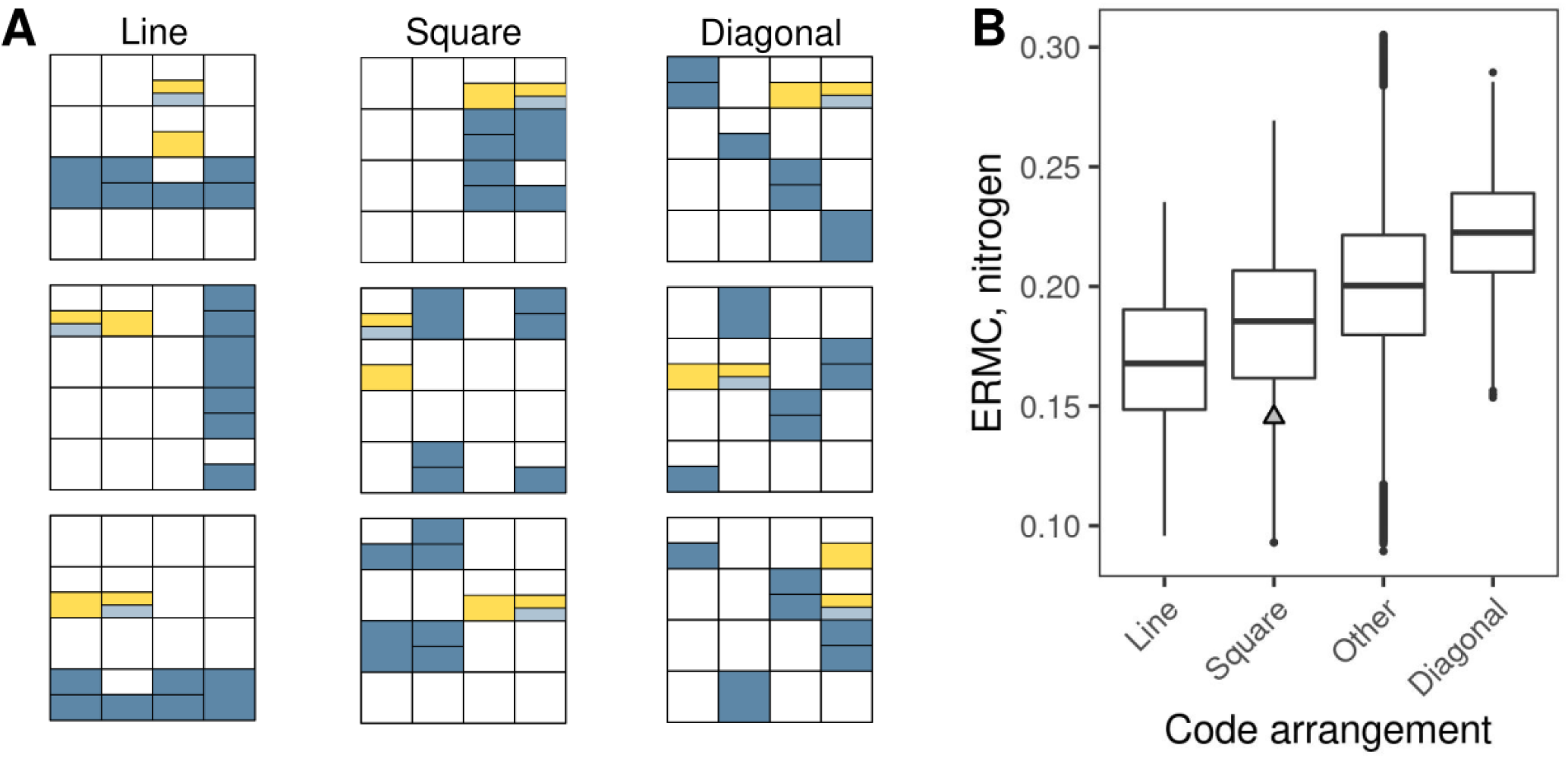
The arrangement of codons for nitrogen-rich amino acids in the codon table influences the ERMC for nitrogen.

A – Schematic depiction of three possible configurations of alternative genetic codes generated by quartet shuffling. Yellow boxes denote stop codons, dark blue boxes denote codons for the amino acids histidine, lysine, asparagine, glutamine, and arginine, and the light blue box is for the tryptophan codon. The topmost square-arranged code is the SGC.
B – The ERMC for nitrogen shown in relation to the different arrangements of codons for nitrogen-rich amino acids in the codon tables of alternative genetic codes, computed using 1 million alternative codes generated by quartet shuffling. The grey triangle corresponds to the SGC. Codes denoted as ‘other’ do not fall into any of the other three categories (‘line’, ‘square’ or ‘diagonal’). The number of codes falling into each of the categories is 5,072 for ‘line’, 20,983 for ‘square’, 961,979 for ‘other’, and 11,966 for ‘diagonal’. The ERMC for nitrogen is computed using the *ocean parameters*.

## Footnotes

1 Massey (2008) instead chooses the subsequent amino acids with equal probability from all amino acids with Grantham distance to the parent amino acid smaller than a chosen threshold; we use an alternative approach to avoid parameterizing the models.

## Notes

### Competing Interest Statement

The authors have declared no competing interest.

## References

Alff-Steinberger C. 1969. The genetic code and error transmission. Proc Natl Acad Sci U S A. 64:584–591.

Archetti M. 2004. Codon usage bias and mutation constraints reduce the level of error minimization of the genetic code. J Mol Evol. 59:258–266.

Ardell DH. 1998. On error minimization in a sequential origin of the standard genetic code. J Mol Evol. 47:1–13.

Athey J, Alexaki A, Osipova E, Rostovtsev A, Santana-Quintero LV, Katneni U, Simonyan V, Kimchi-Sarfaty C. 2017. A new and updated resource for codon usage tables. BMC Bioinformatics. 18:391.

Benjamini Y, Hochberg Y. 1995. Controlling the false discovery rate: A practical and powerful approach to multiple testing. J R Stat Soc Series B Stat Methodol. 57:289–300.

Bragg JG, Wagner A. 2007. Protein carbon content evolves in response to carbon availability and may influence the fate of duplicated genes. Proc Royal Soc B. 274:1063–1070.

Butler T, Goldenfeld N, Mathew D, Luthey-Schulten Z. 2009. Extreme genetic code optimality from a molecular dynamics calculation of amino acid polar requirement. Phys Rev E. 79:060901.

Caporaso JG, Yarus M, Knight RD. 2005. Error minimization and coding triplet/binding site associations are independent features of the canonical genetic code. J Mol Evol. 61:597–607.

Elser JJ, Fagan WF, Subramanian S, Kumar S. 2006. Signatures of ecological resource availability in the animal and plant proteomes. Mol Biol Evol. 23:1946–1951.

Freeland SJ, Hurst LD. 1998. The genetic code is one in a million. J Mol Evol. 47:238–248.

Freeland SJ, Knight RD, Landweber LF, Hurst LD. 2000. Early Fixation of an Optimal Genetic Code. Mol Biol Evol. 17:511–518.

Freeland SJ, Wu T, Keulmann N. 2003. The case for an error minimizing standard genetic code. Orig Life Evol Biosph. 33:457–477.

Gilis D, Massar S, Cerf NJ, Rooman M. 2001. Optimality of the genetic code with respect to protein stability and amino-acid frequencies. Genome Biol. 2:research0049.1.

Goodarzi H, Shateri Najafabadi H, Nejad HA, Torabi N. 2005a. The impact of including tRNA content on the optimality of the genetic code. Bull Math Biol. 67:1355–1368.

Goodarzi H, Shateri Najafabadi H, Torabi N. 2005b. On the coevolution of genes and genetic code. Gene. 362:133–140.

Grantham R. 1974. Amino acid difference formula to help explain protein evolution. Science. 185:862–864.

Grzymski JJ, Dussaq AM. 2012. The significance of nitrogen cost minimization in proteomes of marine microor-ganisms. ISME J. 6:71–80.

Haig D, Hurst LD. 1991. A quantitative measure of error minimization in the genetic code. J Mol Evol. 33:412–417.

Haig D, Hurst LD. 1999. A quantitative measure of error minimization in the genetic code. J Mol Evol. 49:708.

Hellweger FL, Huang Y, Luo H. 2018. Carbon limitation drives GC content evolution of a marine bacterium in an individual-based genome-scale model. ISME J. 12:1180–1187.

Knight RD, Freeland SJ, Landweber LF. 2001. Rewiring the keyboard: evolvability of the genetic code. Nat Rev Genet. 2:49–58.

Li N, Lv J, Niu DK. 2009. Low contents of carbon and nitrogen in highly abundant proteins: Evidence of selection for the economy of atomic composition. J Mol Evol. 68:248–255.

Lv J, Li N, Niu DK. 2008. Association between the availability of environmental resources and the atomic composition of organismal proteomes: Evidence from Prochlorococcus strains living at different depths. Biochem Biophys Res Commun. 375:241–246.

Massey SE. 2008. A neutral origin for error minimization in the genetic code. J Mol Evol. 67:510.

Mazel D, Marlière P. 1989. Adaptive eradication of methionine and cysteine from cyanobacterial light-harvesting proteins. Nature. 341:245–248.

Mende DR, Bryant JA, Aylward FO, Eppley JM, Nielsen T, Karl DM, DeLong EF. 2017. Environmental drivers of a microbial genomic transition zone in the ocean’s interior. Nat Microbiol. 2:1367–1373.

Novozhilov AS, Wolf YI, Koonin EV. 2007. Evolution of the genetic code: partial optimization of a random code for robustness to translation error in a rugged fitness landscape. Biol Direct. 2:24.

Shenhav L, Zeevi D. 2020. Resource conservation manifests in the genetic code. Science. 370:683–687.

Stoltzfus A, Yampolsky LY. 2007. Amino acid exchangeability and the adaptive code hypothesis. J Mol Evol. 65:456–462.

Tripathi S, Deem MW. 2018. The standard genetic code facilitates exploration of the space of functional nucleotide sequences. J Mol Evol. 86:325–339.

Wichmann S, Ardern Z. 2019. Optimality in the standard genetic code is robust with respect to comparison code sets. Biosystems. 185:104023.

Wolf YI, Koonin EV. 2007. On the origin of the translation system and the genetic code in the RNA world by means of natural selection, exaptation, and subfunctionalization. Biol Direct. 2:14.

Xu H, Zhang J. 2021. Unpublished data. https://www.biorxiv.org/content/early/2021/02/16/2021.02.08.430341.full.pdf.

## References

Massey SE. 2006. A sequential “2-1-3” model of genetic code evolution that explains codon constraints. J Mol Evol. 62:809–810.

Massey SE. 2008. A neutral origin for error minimization in the genetic code. J Mol Evol. 67:510.

Plotkin JB, Kudla G. 2011. Synonymous but not the same: the causes and consequences of codon bias. Nat Rev Genet. 12:32–42.

